# Chemotherapy-induced collagen IV drives cancer cell invasion through activation of Src/FAK signaling

**DOI:** 10.1101/2021.04.01.438074

**Authors:** Jackson P. Fatherree, Justinne R. Guarin, Rachel A. McGinn, Stephen P. Naber, Madeleine J. Oudin

## Abstract

Triple-negative breast cancer (TNBC) is the most aggressive and deadly subtype of breast cancer, accounting for 30,000 cases annually in the US. While there are several clinical trials ongoing to identify new agents to treat TNBC, the majority of TNBC patients are treated with anthracycline- or taxane-based chemotherapies in the neoadjuvant setting, followed by surgical resection and adjuvant chemotherapy. While many patients respond well to this approach, as many as 25% will suffer local or metastatic recurrence within five years. Understanding the mechanisms that drive recurrence after chemotherapy treatment is critical to improving survival for patients with TNBC. It is well-established that the extracellular matrix, which provides structure and support to tissues, is a major driver of tumor growth, local invasion and dissemination of cancer cells to distant metastatic sites. In the present study, we show that decellularized extracellular matrix (dECM) obtained from chemotherapy-treated mice increases invasion of treatment-naïve breast cancer cells compared to vehicle-treated dECM. Using tandem-mass-tag proteomics, we further demonstrate that anthracycline- and taxane-based chemotherapies induce drug-specific changes in tumor ECM composition. We identify the basement membrane protein collagen IV as significantly upregulated in the ECM of chemotherapy-treated mice and patients treated with neoadjuvant chemotherapy. We show that collagen IV drives invasion via Src/FAK signaling and that inhibiting collagen IV-driven signaling decreases invasion in chemotherapy-treated dECM. These studies provide a novel mechanism by which chemotherapy may induce metastasis via effects on ECM composition.

**One Sentence Summary:** Chemotherapy alters the extracellular matrix of breast tumors leading to increased invasion of residual cancer cells.

## Main Text

### INTRODUCTION

Triple-negative breast cancer (TNBC) accounts for approximately 15 to 20% of all breast cancers and is associated with high rates of metastasis and poor overall survival *(1)*. This disease is more likely than other subtypes to affect African American and premenopausal women, and represents a significant public health concern *(1)*. While other breast cancer subtypes respond well to therapies targeting hormone receptors (ER/PR) or the receptor tyrosine kinase HER2, TNBC remains extremely difficult to treat due to the lack of targetable driver mutations *(2)*. There are several clinical trials ongoing to identify new agents to treat TNBC. The PD-L1 inhibitor atezolizumab was recently approved for TNBC in combination with paclitaxel, but less than 40% of TNBC patients express PD-L1 and are eligible for this therapy *(3)*. As such, the current standard of care in TNBC is surgical resection followed by cytotoxic chemotherapy, with an increasing number of patients receiving neoadjuvant chemotherapy (NAC) prior to surgery *(4)*. NAC in TNBC consists of a combination of anthracyclines, alkylators and taxanes; the most common regimen consists of four cycles of concurrent doxorubicin (Adriamycin) and cyclophosphamide, followed by four cycles of paclitaxel (Taxol), hereafter referred to as AC-T *(5)*. While NAC increases the likelihood of pathological complete response in TNBC patients, clinical trials have shown that NAC does not improve overall or disease-free survival *(6)*. These clinical observations imply that in a subset of patients, effects on tumor cell proliferation and growth do not correlate with suppression of local invasion which eventually leads to distant metastases.

There is a growing body of evidence indicating that cytotoxic chemotherapy may induce tumor intrinsic and extrinsic changes that promote cancer cell survival and dissemination *(7)*, a phenomenon termed ‘chemotherapy-induced metastasis’. Gene expression analysis of breast cancer patients following NAC reveals an enrichment for highly invasive, stem-like gene signatures in a subset of post-chemotherapy tumors *(8)*. TNBC tumors with low proliferation gene signatures and high cell migration gene signatures were significantly less likely to achieve pathologic complete response to NAC *(9)*. Within the tumor microenvironment (TME), it is well-established that paclitaxel induces tumor angiogenesis by recruiting endothelial progenitor cells *(10, 11)*. Additionally, paclitaxel-induced inflammation promotes the recruitment and outgrowth of myeloid progenitor cells which further support tumor angiogenesis *(12)*. Paclitaxel treatment also stimulates production of epithelial-derived chemoattractants, including colony stimulating factor 1 and interleukin-34, leading to increased recruitment and infiltration of tumor-associated macrophages *(13)*. Doxorubicin also promotes chemotherapy-induced metastasis by inducing vascular leakage leading to recruitment of myeloid-derived cells to the primary tumor *(14)*. Finally, both paclitaxel and doxorubicin have been shown to promote myofibroblastic transformation of fibroblasts in the TME *(15)*. Importantly, many of the stromal cell types implicated in TME-mediated chemotherapy-induced metastasis also dynamically degrade, remodel, and secrete extracellular matrix (ECM). Given the role of the ECM in driving local invasion and metastasis *(16)*, further investigation into the role of tumor ECM in chemotherapy-induced metastasis is warranted.

The ECM, which provides biochemical and biophysical signals to all solid tissues, is a major driver of tumorigenesis and metastasis in breast cancer *(17)*. ECM organization and composition is commonly dysregulated in solid tumors *(18)*. In breast cancer, increased collagen I deposition and fiber alignment is associated with increased invasiveness and high-grade tumors *(19, 20)*. Recent advances in the field of ECM proteomics have revealed exceptional heterogeneity in the tumor matrisome *(21)* and described significant differences in the contributions of cancer cells and stromal cells to ECM deposition *(22)*. It is now understood that the structure and composition of the tumor ECM is the result of dynamic interplay between fibroblasts, macrophages, endothelial cells and cancer cells *(23–26)*. As each of these cell types are directly or indirectly affected by cytotoxic chemotherapy, we hypothesized that chemotherapy would induce substantial changes in the composition of tumor ECM.

Unbiased investigation of the role of ECM in driving cancer cell phenotypes has been an ongoing challenge. *In vitro* investigation of cell-ECM interactions typically involves exposing cells to a single ECM substrate and assaying adhesion, proliferation or invasion. While these tools are useful for dissecting the signals underlying ECM-driven phenotypes, they fail to capture the complexity of native ECM in regard to both composition and 3D structure. More complex tools have been developed to recapitulate native ECM, including the use of Matrigel, cell-derived matrices, and synthetic ECM scaffolds. However, these tools fail to replicate the complexity and heterogeneity of the *in vivo* tumor ECM. To address this, we recently published an experimental pipeline for interrogating the effect of native whole-tissue ECM from different disease states on cancer cells *(27)*. By isolating decellularized ECM (dECM) scaffolds directly from tissues and employing them as a substratum for cell phenotypic studies, we are able to observe whole-tissue ECM-driven cellular phenotypes, irrespective of the cell type that secreted it in the TME.

In the present study, we use this approach to demonstrate a pro-invasive effect of chemotherapy-treated tumor ECM, which is associated with significant alterations in ECM composition. We identify collagen IV as a major driver of TNBC cell invasion that is induced by chemotherapy in both mice and humans. We further show that collagen IV-driven Src/FAK signaling can be targeted to suppress chemotherapy-associated ECM-driven invasion in TNBC. These findings represent the first ECM-based mechanism of chemotherapy-induced metastasis and provide a rationale for targeting Src/FAK signaling in combination with neoadjuvant chemotherapy.

### RESULTS

#### Post-chemotherapy tumor dECM scaffolds drive invasion of TNBC cells

To investigate how cytotoxic chemotherapy alters ECM-driven phenotypes in TNBC, we used transgenic MMTV-PyMT mice, an immunocompetent model that faithfully recapitulates the pathology of human TNBC *(28)*. We grew these mice until their overall tumor burden reached 500 mm^3^, or about 12 weeks of age. At this point, mice were treated every 5 days with 4 cycles of either paclitaxel (10mg/kg), delivered intraperitoneally; doxorubicin (5mg/kg), delivered intravenously; or a vehicle control (Fig. S1A). Both of these treatments significantly slowed tumor growth, but did not induce tumor regression, relative to the vehicle control (Fig. S1B). However, we found no differences in the number of lung metastases at sacrifice between vehicle and chemotherapy-treated mice (Fig. S1C,D).

We then wanted to study the effect of chemotherapy-treated ECM on residual cancer cells. We focused on effects on invasion, which is the first step of metastasis. We decellularized dissected PyMT tumors using our recently published method *(27)* to obtain dECM scaffolds derived from vehicle-, paclitaxel-, or doxorubicin-treated tumors. Decellularization was confirmed by H&E staining (Fig. S2A) and western blot (Fig. S2B-E). These dECM scaffolds were reseeded with GFP-labeled PyMT tumor-derived (PyMT-GFP) cells and imaged live over 16 hours to assay the effect of chemotherapy-treated tumor dECM on 3D invasion speed (Fig. 1A). We found that tumor dECM isolated from either paclitaxel- or doxorubicin-treated tumors significantly increased invasion speed of PyMT-GFP cells relative to vehicle-treated tumors (Fig. 1B-D, Movie S1). These results suggest that chemotherapy-driven changes in the ECM can drive tumor cell invasion, a phenotype that promotes drug resistance, tumor progression and metastasis.

**Fig. 1.**
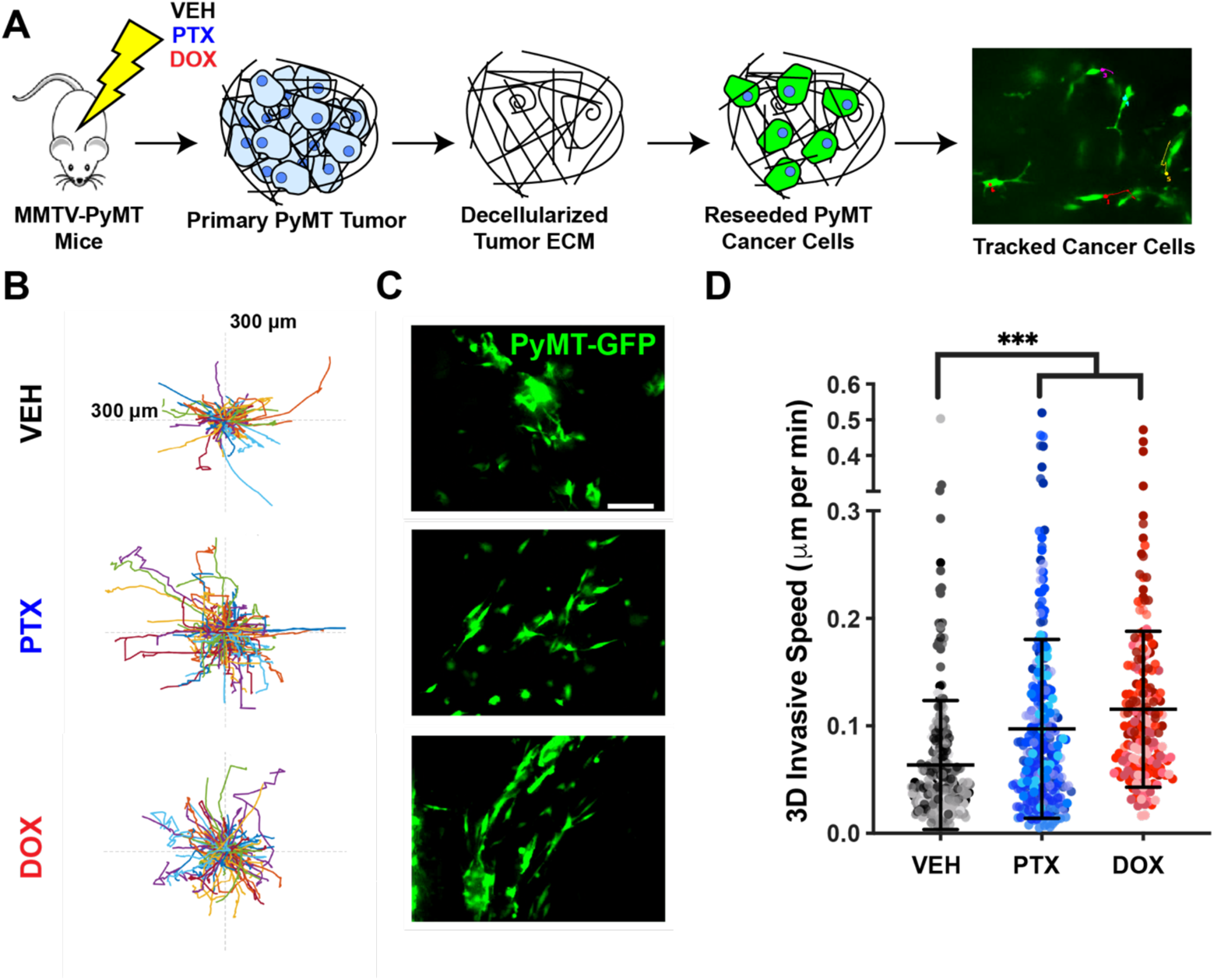
Chemotherapy-treated dECM stimulates invasion of PyMT-GFP TNBC cells. A) Schematic diagram of experiment. MMTV-PyMT mice were treated with paclitaxel (PTX, 10mg/kg), doxorubicin (DOX, 5mg/kg), or a vehicle (VEH) control. Tumors were excised and decellularized to obtain dECM scaffolds. dECM scaffolds were reseeded with PyMT-GFP cells and imaged overnight. Invasive cells were tracked, and invasive speed was quantified. B) Rose plots visualizing the invasive tracks of individual cells on dECM scaffolds. C) Representative frames obtained from reseeding videos of PyMT-cells on dECM from each treatment group (scale bar, 250μm). D) 3D Invasive speed of PyMT-GFP cells seeded on dECM from VEH, PTX and DOX-treated mice. Each point is the average speed of a cell over 16 hours. Like-colored points represent cells tracked on the same dECM scaffold. Data show mean ± SD (n = at least 200 cells tracked from at least 5 dECM scaffolds per treatment group). Significance was determined by one-way ANOVA. \*\*\**P* < 0.005

#### Cytotoxic chemotherapy alters the composition of TNBC tumor ECM

We then sought to investigate how the chemotherapies used here alter the composition of tumor ECM with the goal of identifying protein-level changes that might explain the increased invasive phenotype observed in reseeding experiments. To fully characterize the composition of tumor ECM, we employed a published proteomics approach *(27, 29)*. Existing dECM scaffolds from vehicle- and chemotherapy-treated tumors were first digested to obtain peptides suitable for mass spectrometry analysis. Peptides were then labeled using unique isobaric tandem mass tags (TMTs) to enable quantitative comparison of peptide abundance across samples. We analyzed the resulting dataset using unsupervised hierarchical clustering (Fig. S3) and principal component analysis (PCA), which revealed that paclitaxel and doxorubicin induce distinct changes in the composition of tumor ECM (Fig. 2A). We identified 6 core matrisome proteins in paclitaxel-treated dECM and 17 core matrisome proteins in doxorubicin-treated dECM that were more abundant than in vehicle-treated tumor dECM (log_2_(FC) > 0.3, Fig. 2B,C). These data constitute the first comprehensive characterization of the chemotherapy-treated matrisome. COL4A3 was one of the most significantly upregulated proteins in paclitaxel-treated tumor dECM, so we chose to investigate if chemotherapy treatment induced an increase in native collagen IV deposition in an independent set of MMTV-PyMT mice. We found that both paclitaxel and doxorubicin treatment significantly increase the abundance of native collagen IV in PyMT tumors (Fig. 2D,E). Together, these data indicate that cytotoxic chemotherapy alters the composition of the ECM in tumors and increases collagen IV abundance in the MMTV-PyMT mouse model of TNBC.

**Fig. 2.**
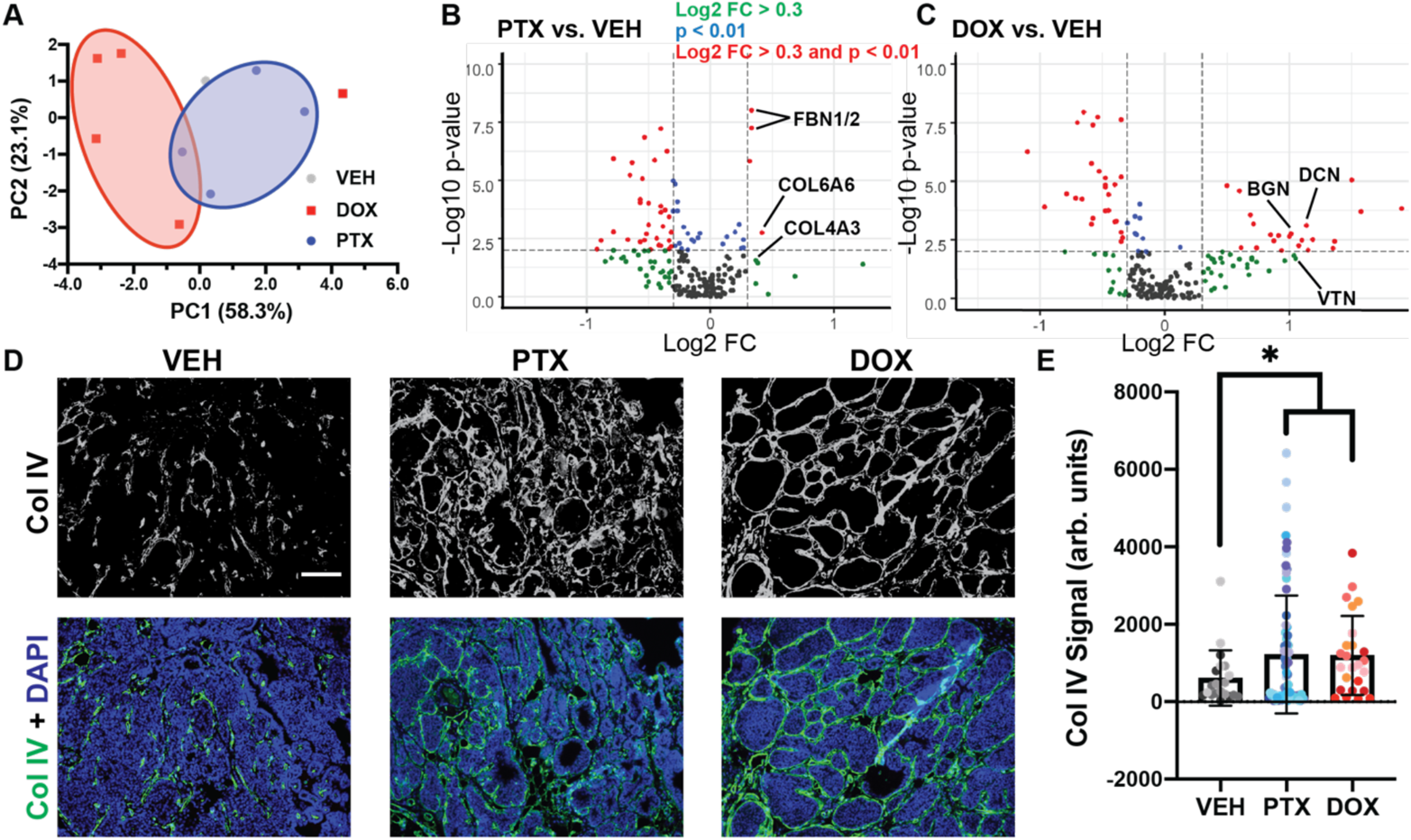
Collagen IV is upregulated by chemotherapy in MMTV-PyMT tumors. A) Principal component analysis of proteomics study investigating dECM composition from VEH, PTX, and DOX-treated tumors. B) Volcano plot comparing dECM composition of PTX- and VEH-treated dECM. C) Volcano plot comparing dECM composition of DOX- and VEH-treated dECM. D) Immunostaining of MMTV-PyMT sections from vehicle- or chemotherapy-treated mice stained for collagen IV and nuclei (scale bar, 100 μm). E) Intensity of collagen IV signal in vehicle and chemotherapy-treated MMTV-PyMT sections. Like-colored points represent fields of view from the same tissue section and mouse. Data show mean ± SD (n = at least 3 dECM scaffolds (A-C) or at least 5 independent mice (D,E). Significance was determined by unpaired *t* test. \**P* < 0.05

#### Treatment with paclitaxel increases collagen IV abundance in human TNBC

We next wanted to determine if collagen IV abundance is also regulated by cytotoxic chemotherapy in human TNBC. We obtained matched diagnostic biopsies and surgical resections from TNBC patients treated at Tufts Medical Center. All patients received AC-T neoadjuvant chemotherapy between diagnosis and surgery. We immunostained sections from diagnostic biopsy (pre-AC-T) and from surgical resections (post-AC-T) to quantify native collagen IV abundance (Fig. 3A). We found that patients displayed a heterogenous range of collagen IV abundance prior to AC-T treatment (Fig. 3B). Interestingly, we found that 3/5 patients displayed an increase in collagen IV abundance greater than 25% (Fig. 3B). These data serve to validate our hypothesis that chemotherapy can induce an increase in collagen IV in human TNBC.

**Fig. 3.**
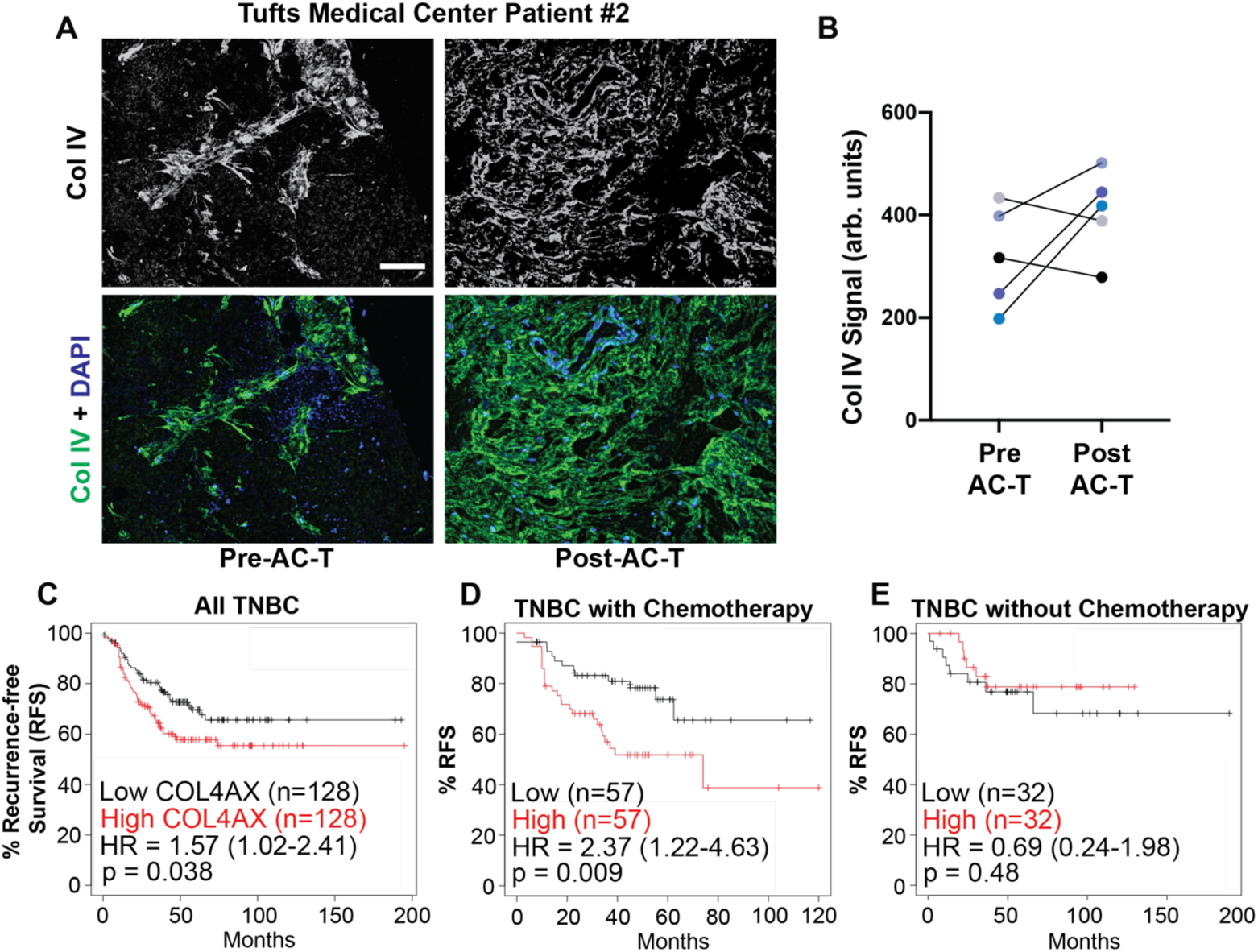
Collagen IV abundance is increased and associated with poor outcomes in chemotherapy-treated human TNBC patients. A) Immunostaining of human TNBC sections retrieved from diagnostic biopsy (pre-AC-T) and surgical resections (post-AC-T) stained for collagen IV and nuclei. (scale bar, 100μm). B) Collagen IV signal intensity measured from matched pre- and post-AC-T sections. C) Kaplan-Meier curve of human TNBC patients based on their average mRNA expression of the six collagen IV α chains (COL4AX). D) Kaplan-Meier curve of human TNBC patients who received chemotherapy comparing low versus high COL4AX expression. E) Kaplan-Meier curve of human TNBC patients who did not receive chemotherapy comparing low versus high COL4AX expression. Patients were stratified by median COL4AX expression and significance was determined by log rank test.

#### Collagen IV is associated with poor outcomes among chemotherapy-treated TNBC patients

To understand the relationship between collagen IV and chemotherapy in a broader clinical context, we interrogated publicly available datasets of TNBC patient outcomes using kmplot.com *(30)*. Collagen IV is encoded by six different genes (COL4A1 to A6), each encoding for a single α chain. These chains organize into 3 heterotrimers (α112, α345, and α556) with distinct tissue distributions. In breast tissue, while α112 is the most common subtype found, all six chains have been detected to various degrees according to the Human Protein Atlas *(31)*. To study clinical outcomes, we stratified patients based on their average expression of all six collagen IV genes (COL4AX). We find that among TNBC patients, high collagen IV expression is associated with poor recurrence-free survival (RFS) (Fig. 3C, HR = 1.57). Interestingly, when we looked deeper into the data, we found that this association was stronger among patients who had received cytotoxic chemotherapy (Fig. 3D, HR = 2.37). Among TNBC patients who did not receive any systemic chemotherapy, there was no association between collagen IV expression and RFS (Fig. 3E, HR = 0.69), further indicating a link between chemotherapy status, collagen IV, and metastatic recurrence. Together, these data demonstrate that cytotoxic chemotherapy increases collagen IV abundance in a subset of TNBC patients, and that this increase may be associated with worse outcomes after chemotherapy treatment.

#### Collagen IV drives 3D invasion of TNBC cells

To determine if increased collagen IV abundance may explain increased invasion observed in cells seeded on chemotherapy-treated dECM, we investigated the effect of collagen IV on the growth and invasion of 3D TNBC spheroids. We previously described a tunable model suitable for analyzing the effect on individual ECM proteins on TNBC cell invasion *(27, 32)*, which accurately replicates the behavior of tumor cells *in vivo (33)*. PyMT-GFP spheroids were embedded in collagen I, the most abundant ECM protein in breast tissue, with or without native human collagen IV and grown for five days. We found that PyMT-GFP spheroids grew significantly larger in the presence of collagen IV when compared to the collagen I only control (Fig. 4A). We also repeated these experiments using GFP-labelled human MDA-MB-231 TNBC cells (231-GFP) and found that collagen IV also increased invasion of these cells (Fig. 4B). Together, these experiments clearly establish that collagen IV stimulates 3D invasion in mouse and human TNBC cells.

**Fig. 4.**
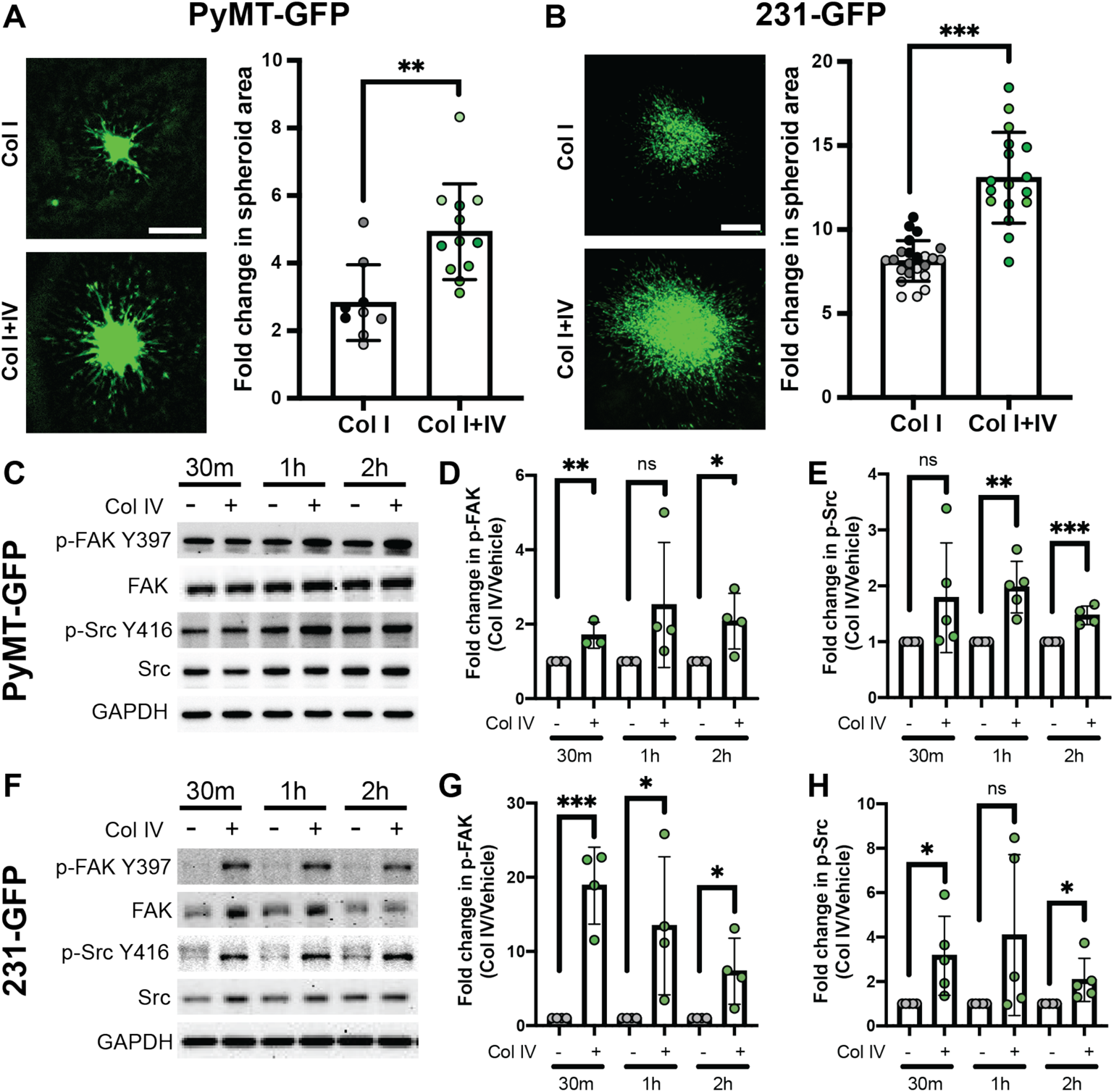
Collagen IV drives 3D invasion and activates Src/FAK signaling in mouse and human TNBC cells. A) Representative images (left) of 3D TNBC spheroids of PyMT-GFP cells grown in collagen I (1mg/mL) with or without 20μg/mL collagen IV (scale bar, 500μm). Fold change in spheroid area was quantified (right). B) 3D TNBC spheroids of 231-GFP cells grown in the same conditions described (scale bar, 500μm). Fold change in spheroid area was quantified (right). Like-colored data points represent technical replicate spheroids grown on the same plate. C-H) Representative western blot images of whole cell PyMT-GFP (C) or 231-GFP (F) lysates immunoblotted for the indicated proteins and phosphoproteins. Cells were plated on collagen IV-coated (Col IV, 20μg/mL) or control dishes for the indicated times before lysis. Quantification of phospho-FAK (Y397) signal in PyMT-GFP (D) or 231-GFP (G) cells relative to total FAK protein and normalized to a loading control. Quantification of phospho-Src (Y416) signal in PyMT-GFP (E) or 231-GFP (H) relative to total Src and normalized to a loading control. Data show mean ± SD (n = at least 9 spheroids from at least 3 independent experiments (A,B) or from at least 3 independent cell lysates (C-H). Significance was determined by unpaired *t* test. \**P* < 0.05, ** *P* < 0.01, *** *P* < 0.005

#### Src/FAK signaling is essential for collagen IV-driven invasion

We next sought to elucidate the molecular pathways underlying collagen IV-driven invasion. Collagen IV has been described to signal through a wide range of pathways depending on the cellular context, including MAPK *(34)*, PI3K *(35)*, Src *(36, 37)*, and FAK *(34, 38)*. To determine which of these signals is mediating collagen IV-driven invasion in our models, we seeded PyMT-GFP or 231-GFP cells on full-length collagen IV-coated dishes for up to 2 hours. Analysis of phosphorylation status of collagen IV-associated signaling pathways revealed activation of Src, as measured by phosphorylation at Y416, and FAK, as measured by phosphorylation at Y397, in both PyMT-GFP and 231-GFP cells (Fig. 4C-H). We found no activation of MAPK or PI3K pathways as measured by Erk1/2 phosphorylation at T202/204 and Akt phosphorylation at S473 respectively (Fig. S4).

To investigate the importance of these pathways in collagen IV-driven 3D invasion, we leveraged existing small molecule inhibitors targeting Src (dasatinib) and FAK (defactinib). Using our spheroid model of 3D invasion, PyMT-GFP or 231-GFP cells were seeded with or without collagen IV and treated with 10nM dasatinib or 1μM defactinib. These doses were chosen to detect anti-invasive effects while having minimal effect on cell proliferation or viability as described previously *(39–41)*. In PyMT-GFP cells, inhibition of FAK, but not Src, reduced 3D invasion in collagen I, in the absence of collagen IV. However, inhibition of either Src or FAK reduced the collagen IV-driven increase in 3D invasion (Fig. 5A,B). Further, 231-GFP cells were insensitive to both Src and FAK inhibition in the absence of collagen IV. Src or FAK inhibition ablated collagen IV-stimulated invasion in these cells as well (Fig. 5C,D), demonstrating the importance of Src and FAK signaling in collagen IV-driven TNBC invasion.

**Fig. 5.**
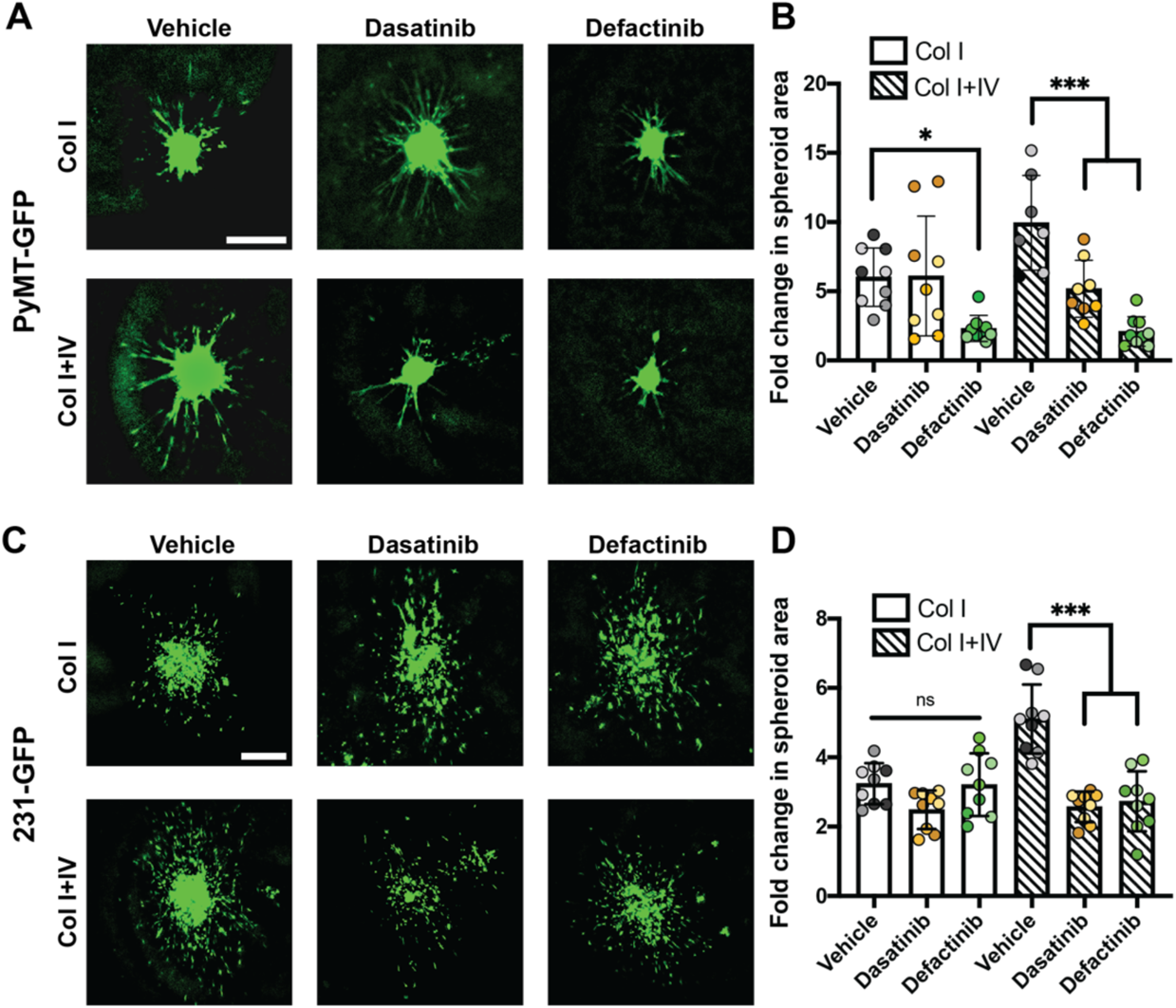
Src or FAK inhibition ablates collagen IV-driven invasion in TNBC cells. A) Representative images of PyMT-GFP spheroids grown with or without collagen IV and treated with DMSO, 10nM dasatinib or 1μM defactinib (scale bar, 500μm). B) Quantification of fold change in PyMT-GFP spheroid area after four days. Each dot represents an individual spheroid. C) Representative images of 231-GFP spheroids grown with or without collagen IV and treated with DMSO, 10nM dasatinib or 1μM defactinib (scale bar, 500μm). D) Quantification of fold change in MDA-MB-231 spheroid area after four days. Each dot represents an individual spheroid. Like-colored data points represent technical replicate spheroids grown on the same plate. Data show mean ± SD (n= 7-9 spheroids pooled from 3 independent experiments). Significance was determined by one-way ANOVA. \**P* < 0.05, \*\*\**P* < 0.005

#### Blocking collagen IV signaling suppresses chemotherapy-associated dECM-driven invasion

We last sought to confirm that collagen IV-driven signaling accounts for the increased invasion induced by chemotherapy-treated dECM scaffolds identified in Fig 1C. PyMT-GFP cells were seeded onto paclitaxel- and vehicle-treated dECM scaffolds and treated with 10nM dasatinib or 1uM defactinib, consistent with the doses found to suppress collagen IV-driven invasion in our spheroid assays (Fig 5). We found that neither drug suppressed invasion of cells seeded on vehicle-treated dECM (Fig. 6A,B, Movie S2). Either drug alone effectively suppressed chemotherapy-associated invasion of cells seeded on paclitaxel-treated dECM (Fig. 6A,C, Movie S3). To determine if chemotherapy-associated ECM-driven invasion was under the control of integrin β1 as previously reported *(34, 42)*, we treated reseeded PyMT-GFP cells with an integrin β1 blocking antibody. Surprisingly, this treatment had no effect on invasion on either paclitaxel- or vehicle-treated dECM (Fig. 6B,C), implying the presence of other integrin or non-integrin receptors for transducing signals from collagen IV into the cell. We then seeded 231-GFP human TNBC cells onto vehicle- or paclitaxel-treated dECM and subjected them to Src or FAK inhibition. Consistent with our previous data, we found that targeting Src or FAK had no effect on invasion on vehicle-treated dECM (Fig. 6D,E, Movie S4), but completely suppressed chemotherapy-associated invasion on paclitaxel-treated dECM (Fig. 6D,F, Movie S5). Together, these data demonstrate that collagen IV-associated signaling potentiates ECM-driven invasion in post-chemotherapy dECM scaffolds.

**Fig. 6.**
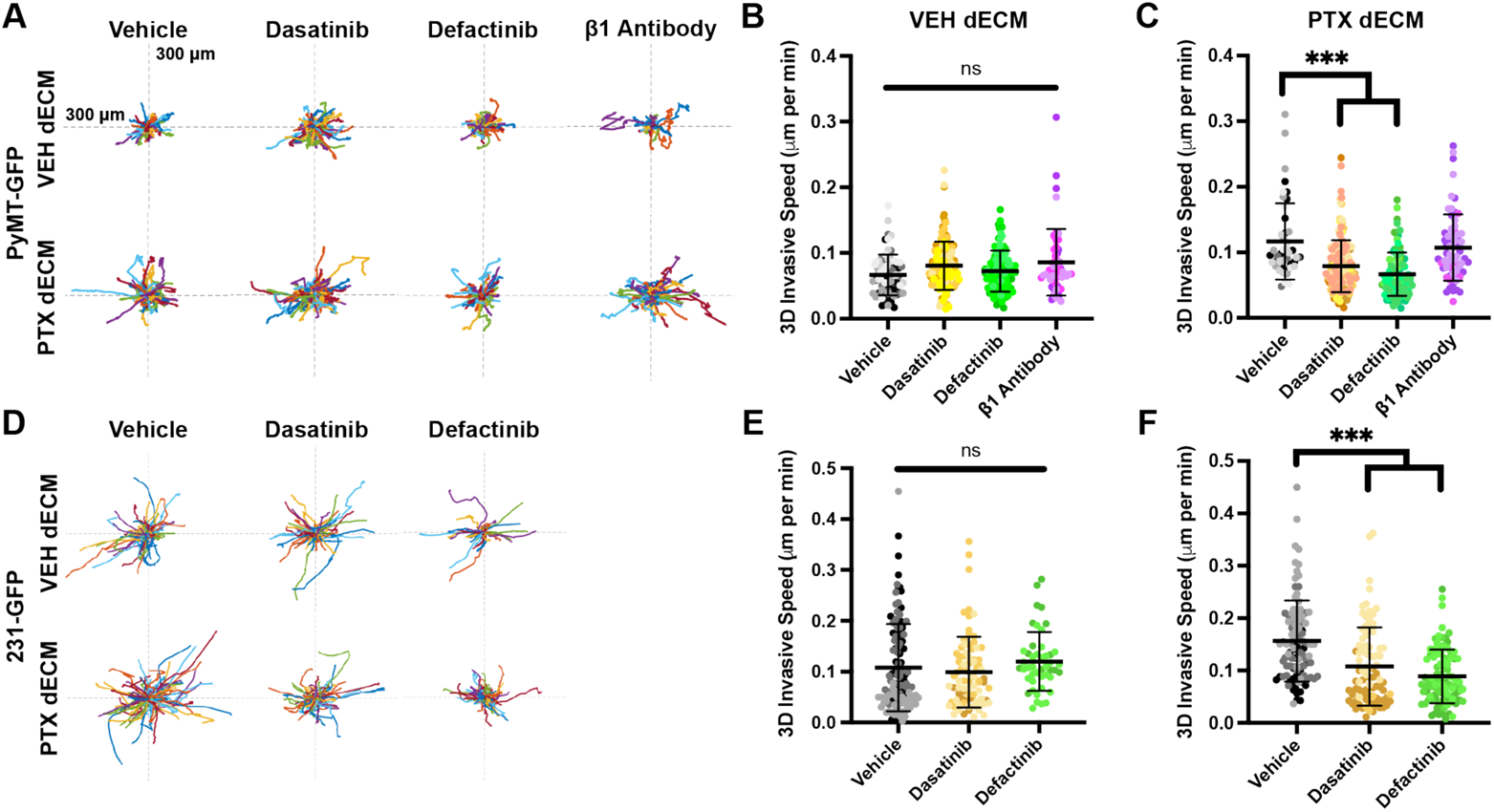
Inhibition of Src or FAK, but not integrin β1, suppresses chemotherapy-associated ECM-driven invasion of TNBC cells. A) Rose plots depicting invasive tracks of individual PyMT-GFP cells seeded onto vehicle- or paclitaxel-treated dECM scaffolds and treated with DMSO, dasatinib (10nM), defactinib (1μM), or an integrin β1 blocking antibody (100ng/mL). B) Quantification of invasive speed of PyMT-GFP cells seeded on vehicle-treated dECM. C) Quantification of invasive speed of PyMT-GFP cells seeded on paclitaxel-treated dECM. D) Rose plots depicting invasive tracks of individual MDA-MB-231 cells seeded onto vehicle- or paclitaxel-treated dECM scaffolds and treated with DMSO, dasatinib (10nM), or defactinib (1μM). E) Quantification of invasive speed of MDA-MB-231 cells seeded on vehicle-treated dECM. F) Quantification of invasive speed of MDA-MB-231 cells seeded on paclitaxel-treated dECM. Each dot represents an individual tracked cell. Like-colored data points represent cell tracked from the same dECM scaffold. Data show mean ± SD (n = at least 45 cells tracked from at least 3 dECM scaffolds per treatment group). Significance was determined by one-way ANOVA. \*\*\**P* < 0.005

### DISCUSSION

Cytotoxic chemotherapy is an important tool that benefits many thousands of patients each year across cancer types. However, resistance to therapy and recurrence post-treatment occurs in over 25% of TNBC patients. Therefore, it is essential that we gain a better understanding of the breadth of the systemic changes caused by these therapies to maximize their clinical benefit and identify new clinical approaches to reduce the risk of chemotherapy-induced metastasis. While it is now understood that the ECM is an essential driver of tumor cell survival, invasion, and metastasis, the effect of chemotherapy on the ECM had not been investigated. Here, we report the first characterization of the chemotherapy-treated matrisome and identify collagen IV as a driver of invasion in the post-chemotherapy tumor microenvironment. Importantly, we found that collagen IV expression was associated with increased likelihood of metastasis only in TNBC patients treated with cytotoxic chemotherapy, indicating that chemotherapy-induced collagen IV may be particularly conducive to TNBC metastasis. Using our dECM model, we further demonstrated that collagen IV-induced Src and FAK signaling were essential to cancer cell invasion only in the context of chemotherapy-treated, collagen IV-rich dECM. Together, these data illustrate the first ECM-driven mechanism of chemotherapy-induced metastasis identified in TNBC.

Collagen IV is an essential basement membrane protein found in all tissues of the body that plays a variety of roles in tumor progression. While it is not found to be upregulated in primary breast cancer *(43)*, serological collagen IV abundance positively correlates with breast cancer staging *(44)* and functions as a prognostic marker of breast cancer metastasis *(45)*. It has been shown to stimulate adhesion and invasion in a wide range of cancer types *(34, 35, 46)*, however the signaling pathways required for collagen IV-driven invasion in TNBC have not been identified. Here, we report that co-activation of Src and FAK is essential for collagen IV-driven invasion in 3D. Further, we demonstrate a novel role of collagen IV in driving cancer cell invasion in the post-chemotherapy tumor microenvironment. We show here that blockade of integrin β1 does not suppress cancer cell invasion in post-chemotherapy dECM, but we were unable to determine the tumor cell receptor responsible for transducing chemotherapy-associated collagen IV-driven signals. Collagen IV has been shown to stimulate invasion through a number of other receptors, including integrin αvβ3 and DDR1 *(47, 48)*. Further study is required to determine if targeting these receptors can inhibit chemotherapy-associated ECM-driven invasion.

The present study raises a number of questions about the microenvironmental response to NAC in TNBC. First, the mechanism by which chemotherapy stimulates collagen IV production *in vivo* is unclear. Proteomics studies have established that collagen IV is produced in similar amounts by both tumor and stromal cells *in vivo (29)*. In the stromal compartment, fibroblasts, macrophages and endothelial cells have all been shown to produce collagen IV *(49–51)*. Cui *et al.* found that cancer-associated fibroblasts produce collagen IV in response to taxane-based chemotherapy *in vitro*, but this has not been observed *in vivo (49)*. Chemotherapy stimulates secretion of tumor-derived soluble signals leading to paracrine crosstalk between tumor and stromal cells *(52)*, which could be another potential mechanism driving chemotherapy-induced ECM production. Further, our proteomics experiment identified a number of potentially pro-metastatic ECM proteins associated with chemotherapy that can serve as targets for future investigation. Paclitaxel treatment was associated with increased collagen VI abundance, which we have recently shown drives invasion in TNBC *(27)*. Fibrillin-1 and -2 were also identified in paclitaxel-treated dECM. While understudied in breast cancer, these proteins have pro-tumor effects in ovarian and lung cancer respectively *(53, 54)*. In doxorubicin-treated dECM, we identified vitronectin and biglycan, two proteins known to promote invasiveness in cancer *(55, 56)*. Interestingly, we also identified the tumor suppressive protein decorin *(57)*. Further study is required to fully characterize the effect of these complex chemotherapy-induced ECM changes on cancer cell phenotypes. While the present study focused on the compositional changes in ECM observed in the tumors, structure, crosslinking and physical properties of the ECM can also promote tumor progression *(16)*. Therefore, future studies are needed to evaluate the effect of chemotherapeutic drugs on the production of ECM or chemokines that may stimulate ECM production, by cells in the TME, as well as on the effect of chemotherapy on the biophysical properties of the ECM.

Overall, our work reveals a clinical paradigm where an increase in collagen IV induced by NAC may predispose patients to suffer metastatic recurrence, which could be translated to patients in several ways. First, given that tumor-derived collagen IV can be detected in the serum of breast cancer patients *(44)*, it is possible measuring that collagen IV could function as a biomarker in the blood to non-invasively track drug response and predict disease progression. Further, the ECM is a promising target to facilitate drug delivery. Collagen-mimetic peptides have been developed that enable delivery of cargo to locations of high ECM turnover in the body, including to solid tumors *(58)*. ECM-targeting antibodies have been developed that facilitate cargo delivery to tissues rich in specific ECM proteins *(59)*. This technology could be used in patients with high collagen IV following chemotherapy to deliver metastasis-suppressing drugs. Finally, this study employed two FDA-approved small molecule inhibitors targeting Src (dasatinib) and FAK (defactinib). Our data suggests that these drugs may suppress cancer cell dissemination in early stage TNBC, allowing for adjuvant chemotherapy to more effectively clear residual cancer cells. Dasatinib has been employed in combination with paclitaxel in breast cancer and has shown to be well-tolerated *(60)*. However, clinical trials combining these drugs have been limited to metastatic breast cancer patients *(61, 62)*. In one phase II study, 43% of patients showed either tumor regression or stable disease for greater than 6 months, but was discontinued due to slow patient accrual *(61)*. While multiple phase I studies have demonstrated safety of defactinib in patients *(63, 64)*, no completed studies have tested the efficacy of defactinib in combination with chemotherapy in breast cancer. The present study builds on existing preclinical literature demonstrating the efficacy of defactinib in overcoming chemoresistance *(65)*, suppressing metastasis *(66)*, and eliminating cancer stem cells *(67)* in a range of solid tumors. Taken together, our studies shed light on novel mechanisms of chemotherapy-induced metastasis in TNBC involving the ECM, that have the potential for clinical translation and development of new strategies to track, predict and overcome drug resistance to prevent further metastasis and to reduce patient death.

### MATERIALS AND METHODS

#### Study Design

This study was designed to investigate ECM-driven phenotypes associated with chemotherapy treatment in TNBC. Specifically, we sought to use our novel dECM pipeline to characterize how chemotherapy changes tumor ECM composition, and how those changes affect cancer cell behavior. We used the immunocompetent MMTV-PyMT genetic mouse model of TNBC to ensure detection of chemotherapy-induced ECM remodeling regardless of the driving cell type (fibroblast, endothelial cell, tumor cell, etc.). Mice were randomized into treatment groups upon reaching a tumor burden of 500 mm^3^ and were only removed from the study if they reached their tumor burden endpoint (any tumor exceeding 20 mm in one direction) prior to cessation of treatment. Paclitaxel and doxorubicin were chosen as chemotherapeutics to test as they are employed in AC-T therapy, the most common chemotherapy regimen used in the neoadjuvant setting. We determined that a sample size of 12 mice per treatment group was sufficient based on a power analysis with the assumption that the difference between groups would be greater than 1.5 times the within-group standard deviation (two-tailed alpha of 0.5).

Discovery and validation of collagen IV as a chemotherapy-induced ECM protein was conducted using tissues from separate mice to reduce the chance that this observation was an artifact of a single experiment. Researchers were blinded during imaging and quantification of collagen IV-stained MMTV-PyMT tissue sections. Human TNBC sections from diagnostic biopsy and surgical resection are easily distinguishable due to size and density of tissue section so researchers were not blinded from those experiments. *In vitro* experiments in this study were repeated a minimum of three times to ensure reproducibility. We excluded data from reseeding assays only when it was apparent that cells were not adhered to dECM, such as when the majority of cells in the field of view died during the experiment. All other data that was produced in the discussed experiments was presented.

#### Mice

All animal studies were reviewed and approved by the Tufts University Institutional Animal Care and Use Committee. Transgenic mice bearing the polyomavirus middle T antigen under control of the mouse mammary tumor virus promoter (MMTV-PyMT) were obtained from The Jackson Laboratory (Bar Harbor, ME). MMTV-PyMT mice were grown until overall tumor burden reached 500 mm^3^ at about 10-12 weeks of age before being randomized into chemotherapy treatment groups. Paclitaxel (Taxol) was resuspended in 5% dimethyl sulfoxide, 40% polyethylene glycol 3000, and 5% Tween 80 in dH_2_O and administered intraperitoneally at 10 mg/kg. Doxorubicin (Adriamycin) was dissolved in phosphate-buffered saline and administered intravenously at 5 mg/kg. Appropriate vehicle controls for each drug were included in the control group. All drugs were administered for 4 cycles given every 5 days. Tumor burden was monitored using digital calipers at each treatment. Three days after the final treatment, mice were euthanized, and tumors and lungs were excised for further study.

#### Antibodies and inhibitors

Primary antibodies used in this study include: α-fibronectin (ab2413, Abcam, Cambridge, MA), α-tubulin (DM1A, Sigma-Aldrich, St. Louis, MO), α-GAPDH (14C10, Cell Signaling Technology, Danvers, MA), α-collagen IV (ab6586, Abcam), α-phospho-Src (2101, Cell Signaling Technology), α-Src (36D10, Cell Signaling Technology), α-phospho-FAK (3283, Cell Signaling Technology), α-FAK (D5O7U, Cell Signaling Technology), α-phospho-Erk1/2 (4370, Cell Signaling Technology), α-Erk1/2 (4695, Cell Signaling Technology), α-phospho-Akt (9271, Cell Signaling Technology), α-Akt (4691, Cell Signaling Technology) and α-integrin β1 (1:100, P4C10, Millipore, Burlington, MA). All antibodies were used at a concentration of 1:1000 unless indicated otherwise. Pharmacological inhibitors include: dasatinib (S1021, Selleck Chemicals, Houston, TX) and defactinib (S7654, Selleck Chemicals).

#### Cell Culture

231-GFP cells were obtained from American Type Cell Collection (Mannassas, VA) and cultured in Dulbecco’s modified Eagle’s medium (DMEM) with 10% fetal bovine serum and 1% penicillin-streptomycin-glutamine. PyMT-GFP cells were a gift Prof. Richard Hynes’ lab at MIT and were cultured in a 1:1 mix of DMEM and Ham’s F-12 Nutrient Mixture containing 2% fetal bovine serum, 1% bovine serum albumin (A2153-50G; Sigma-Aldrich, St. Louis, MO), EGF (10 ng/ml; PHG0313; Fisher Scientific, Hampton, NH), insulin (10 μg/ml; 12585014; Fisher Scientific, Hampton, NH), and 1% penicillin-streptomycin-glutamine. Cells were monitored for mycoplasma contamination by polymerase chain reaction (PCR) using the Universal Mycoplasma Detection Kit (30-1012 K; ATCC, Manassas, VA). Only mycoplasma-negative cells were used in this study.

#### Isolation of dECM scaffolds

Tumor-derived dECM scaffolds were produced as described previously *(27)*. Briefly, tumors dissected from MMTV-PyMT mice were submerged in 0.1% SDS in PBS (w/v) solution with agitation for several days, replacing the solution twice daily. Upon successful decellularization, tissues were moved to 0.05% Triton X-100 in PBS solution for 3 hours, followed by washing in dH_2_O for 48 hours to remove residual detergents. To assess decellularization, dECM scaffolds were paraffin embedded, sectioned and H&E stained.

#### Protein Isolation

dECM, intact tumors, or cell pellets were lysed in 25 mM tris, 150 mM NaCl, 10% glycerol, 1% NP-40, and 0.5 M EDTA with 1× protease Mini-complete protease inhibitor (04693124001; Roche, Indianapolis, IN) and 1× phosphatase inhibitor cocktail (4906845001; Roche, Indianapolis, IN) at 4°C. In the case of dECM and intact tumors, mechanical homogenization was achieved using the Bead-Bug 3 Place Microtube Homogenizer (D1030; Benchmark Scientific, Sayreville, NJ). The resulting homogenate was then centrifuged at 21,000g for 10 minutes at 4C and the supernatant was stored at −20C for subsequent analysis.

#### dECM Reseeding Experiments

Reseeding experiments were performed as described previously *(27)*. 5mm pieces of dECM scaffold were cut and conditioned in complete cell culture media for 48 hours. On the morning of the experiment, tissues were reseeded with 250,000 231-GFP or PyMT-GFP cells and incubated at 37C for 6 hours. Reseeded tissue was then moved to fresh media, stabilized, and imaged for 16 hours, capturing images every 20 minutes. For inhibitor experiments, vehicle controls, antibodies, or small-molecule inhibitors were added to the media just before imaging began. Cells were then tracked using VW-9000 Video Editing/Analysis Software (Keyence, Elmwood Park, NJ), and invasive speed was calculated using a custom MATLAB script vR2018a (MathWorks, Natick, MA). Data presented are the result of at least three independent experiments with five fields of view imaged per experiment and 4-8 cells tracked per field of view.

#### LC-MS/MS Analysis

Samples were prepared as described previously *(27)*. Briefly, dECM homogenate was denatured in 8M urea and 10mM dithiothreitol, alkylated with 25mM iodoacetamide, and deglycosylated with peptide *N*-glycosidase F (P0704S; New England Biolabs, Ipswich, MA). Samples were then digested sequentially, first with endoproteinase LysC (125-05061; Wako Chemicals USA, Richmond, VA), then trypsin (PR-V5113; Promega, Madison, WI). Samples were acidified with 50% trifluoroacetic acid and labeled with TMT10plex (90110; Thermo Fisher Scientific, Waltham, MA) according to the manufacturer’s instructions. Labeled peptides were fractionated via high-pH reverse-phase HPLC (Thermo Easy nLC1000, Thermo Fisher Scientific, Waltham, MA) using a precolumn (made in house, 6 cm of 10-μm C18) and a self-pack 5-μm tip analytical column (12 cm of 5-μm C18; New Objective, Woburn, MA) over a 140-min gradient before nanoelectrospray using a Q Exactive HF-X mass spectrometer (Thermo Fisher Scientific, Waltham, MA). Solvent A was 0.1% formic acid, and solvent B was 80% ACN/0.1% formic acid. The gradient conditions were 2 to 10% B (0 to 3 min), 10 to 30% B (3 to 107 min), 30 to 40% B (107 to 121 min), 40 to 60% B (121 to 126 min), 60 to 100% B (126 to 127 min), 100% B (127 to 137 min), 100 to 0% B (137 to 138 min), and 0% B (138 to 140 min), and the mass spectrometer was operated in a data-dependent mode. The parameters for the full-scan MS were resolution of 60,000 across 350 to 2000 m/z (mass/charge ratio), automatic gain control 3 x 106, and maximum ion injection time of 50 ms. The full-scan MS was followed by tandem mass spectrometry (MS/MS) for the top 15 precursor ions in each cycle with a normalized collision energy of 34 and dynamic exclusion of 30 s.

#### Western Blotting

Protein lysates were separated using SDS-polyacrylamide gel electrophoresis on a 12% polyacrylamide gel. Proteins were transferred to a nitrocellulose membrane using a TransBlot Turbo Transfer system (Bio-Rad; Hercules, CA). Membranes were blocked in 5% nonfat dry milk in tris-buffered saline with 0.05% tween 20 and incubated in primary antibody overnight at 4C with rocking. Antibody binding was visualized using horseradish peroxidase-conjugated secondary antibodies (Jackson ImmunoResearch, West Grove, PA). Imaging was performed using a ChemiDoc MP imaging system (12003154; Bio-Rad, Hercules, CA).

#### Immunohistochemistry

Tissue fixation, processing and sectioning was performed as previously described *(27)*. MMTV-PyMT tumors were fixed in 4% paraformaldehyde in PBS, embedded in paraffin, and sectioned into 10μm sections. Human TNBC samples were obtained from the Tufts Medical Center Biorepository as deidentified formalin-fixed, paraffin-embedded 5μm sections. For H&E staining, sections were deparaffinized, hydrated, stained with hematoxylin (GHS280; Sigma-Aldrich, St. Louis, MO), and counterstained with eosin (HT110180; Sigma-Aldrich, St. Louis, MO). Stained sections were mounted in toluene (SP15-500, Fisher Scientific, Hampton, NH). For immunofluorescence, tissue sections were deparaffinized and antigen retrieval was performed in Citra Plus solution (HK057; Biogenex, Fremont, CA). Sections were then blocked in PBS with 0.5% tween 20 and 10% donkey serum and incubated with primary antibodies overnight at 4C. The next day, sections were incubated with fluorophore-conjugated secondary antibodies and 4′,6-Diamidino-2-phenylindole (DAPI; D1306; Thermo Fisher Scientific, Waltham, MA) to stain cell nuclei. Sections were mounted in Fluoromount mounting medium (00-4958-02; Thermo Fisher Scientific, Waltham, MA) and imaged using a Keyence BZ-X710 microscope (Keyence, Elmwood Park, NJ). Quantification of ECM signal was performed using ImageJ (National Institutes of Health, Bethesda, MD).

#### Spheroid Invasion Assay

Spheroid invasion assays were performed as described previously *(32)*. 2,000 231-GFP or PyMT-GFP cells were seeded in round-bottom, low-attachment 96-well plates and centrifuged at 3,000g for 3 minutes to induce spheroid formation. Spheroids were allowed to grow for three days before addition of an ECM mixture of 1mg/mL collagen I (354236; Corning, Corning, NY), 10mM NaOH, 7.5% 10X DMEM, and 50% DMEM with or without 20μg/mL native human collagen IV (ab7536; Abcam, Cambridge, MA). Once ECM had solidified, 50uL of culture media to maintain moisture and humidity in the well. In inhibitor studies, this media included DMSO as a vehicle control or small molecule inhibitors. Spheroids were imaged in 3D on the day of ECM addition and after 4 days of growth using a Keyence BZ-X710 microscope (Keyence, Elmwood Park, NJ). Data presented are the result of three independent experiments with three technical replicates per experiment.

#### Statistical Analysis

Statistical analysis and visualization were performed using GraphPad Prism v8.4.3. To compare two groups, an unpaired Student’s *t* test was used where a P value ≤ 0.05 was considered statistically significant. To compare between more than two groups, a one-way analysis of variance (ANOVA) was performed using a Bonferroni multiple testing correction where a corrected P value ≤ 0.05 was considered statistically significant. For analysis of patient survival, a log rank test was used where a P value ≤ 0.05 was considered statistically significant.

## Supplementary Materials

Fig. S1. Cytotoxic chemotherapy slows tumor growth but not pulmonary metastasis.

Fig. S2. Decellularization removes cellular material while preserving ECM components.

Movie S1. PyMT-GFP cells seeded onto vehicle-, paclitaxel-, or doxorubicin-treated dECM scaffolds.

Fig. S3. Paclitaxel and doxorubicin treatment induce largely distinct changes in tumor ECM composition.

Table S1. Results of proteomics study comparing chemotherapy- and vehicle-treated dECM scaffolds.

Fig. S4. MAPK and PI3K pathways are not activated by collagen IV in PyMT-GFP TNBC cells.

Movie S2. PyMT-GFP cells seeded on vehicle-treated dECM scaffolds are insensitive to Src, FAK, and integrin β1 inhibition.

Movie S3. Inhibition of Src or FAK, but not integrin β1, suppresses PyMT-GFP invasion on paclitaxel-treated dECM.

Movie S4. 231-GFP cells seeded on vehicle-treated dECM scaffolds are insensitive to Src or FAK inhibition.

Movie S5. Inhibition of Src or FAK suppresses 231-GFP invasion on paclitaxel-treated dECM.

## Funding

This work was supported by:

National Institutes of Health [R00-CA207866 to M.J.O.]

Tufts University [Start-up funds from the School of Engineering to M.J.O.],

Tufts Graduate School of Biomedical Sciences [Collaborative Cancer Biology Award to J.P.F.]

## Author contributions

Conceptualization: JPF, MJO

Methodology: JPF, JRG, SN, MJO

Investigation: JPF, JRG, RAM,SN

Visualization: JPF, JRG, RAM,SN

Funding acquisition: JPF, MJO

Project administration: JPF, SN, MJO

Supervision: JPF, MJO

Writing – original draft: JPF, MJO

Writing – review & editing: JPF, JRG, RAM, SN, MJO

## Data and materials availability

All data are available in the main text or the supplemental materials.

## Supplementary materials

**Fig. S1.**
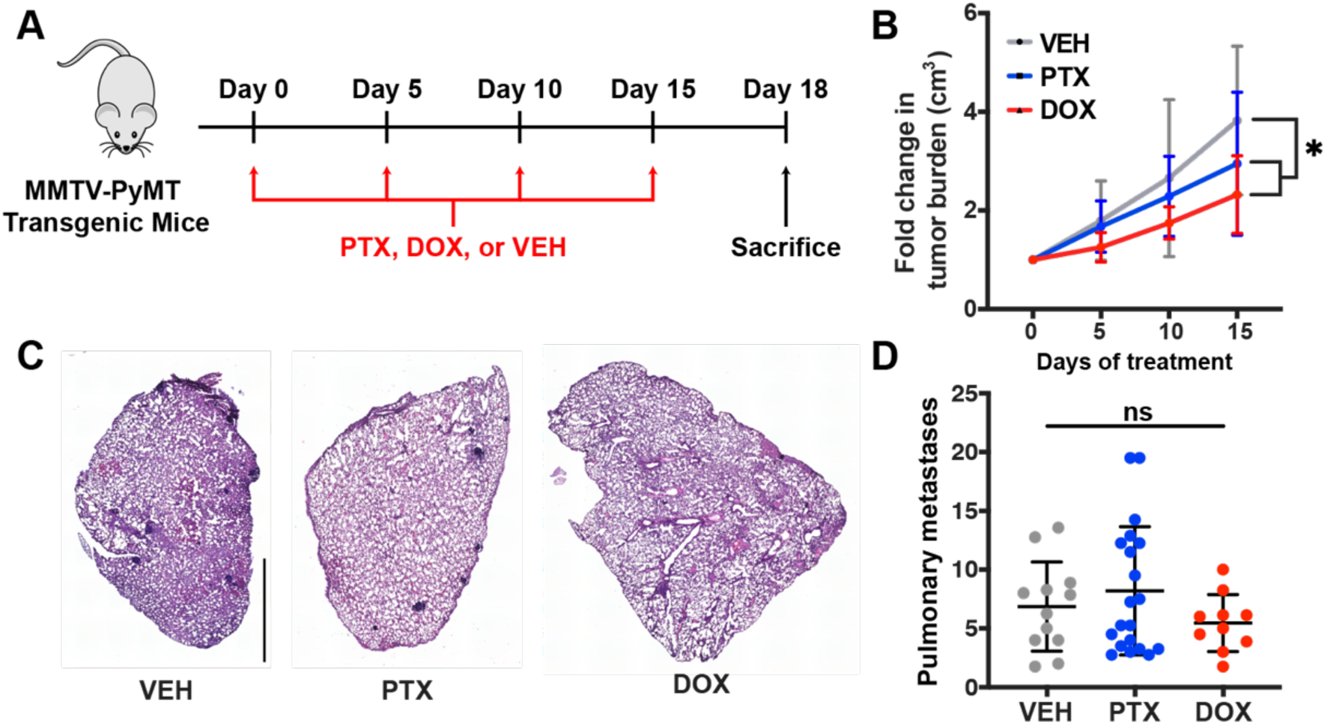
Cytotoxic chemotherapy slows primary tumor growth but not pulmonary metastasis in MMTV-PyMT mice. A) Schematic of *in vivo* approach. MMTV-PyMT mice were treated with four cycles of paclitaxel (PTX, 10mg/kg), doxorubicin (DOX, 5mg/kg), or a vehicle control, delivered every five days. Three days after the final treatment mice were euthanized for downstream analysis. B) Fold change in tumor burden over the course of the experiment. Both paclitaxel and doxorubicin slow tumor growth compared to a vehicle control. C) Representative H&E images of lung sections (scale bar, 1mm). D) Quantification of pulmonary metastases per mouse lung averaged from multiple independent and blinded counts. Data show mean ± SD (n = at least 10 mice per treatment group). Significance was determined by one-way ANOVA. \**P* < 0.05

**Fig. S2.**
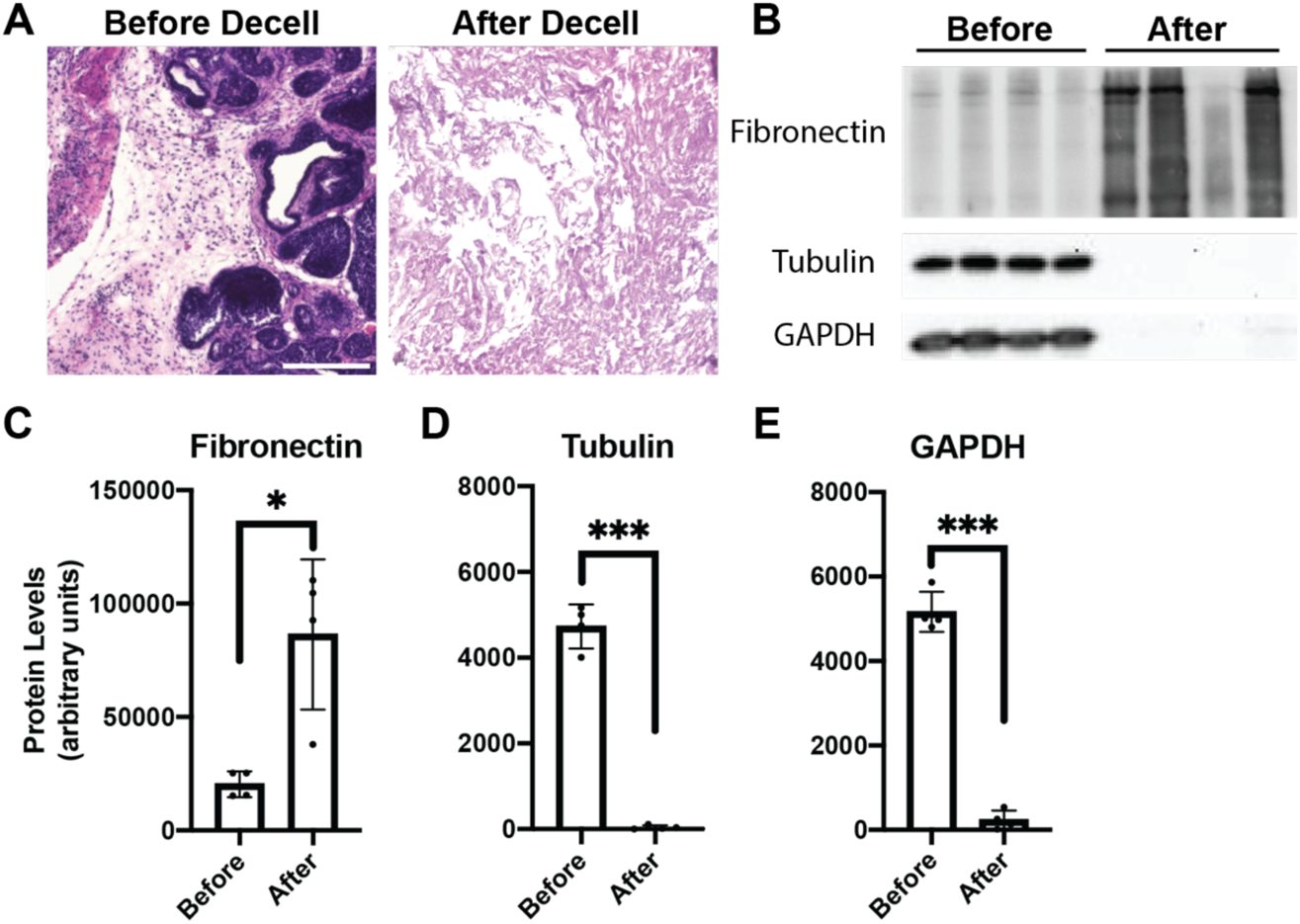
Decellularization removes cellular material while preserving ECM components. A) Representative H&E images of MMTV-PyMT tumor sections before and after decellularization (scale bar, 100μm). B) Western blot analysis of protein lysates extracted from frozen MMTV-PyMT tumors (Before) or dECM scaffolds (After). Fibronectin blot demonstrates retention of ECM whereas tubulin and GAPDH blots demonstrate removal of cellular material. C-E) Quantification of signal intensity in fibronectin (C), tubulin (D), and GAPDH (E) blots. Data show mean ± SD (n = 4 independent tumors or dECM scaffolds). Significance was determined by unpaired *t* test. * *P* < 0.05, *** *P* < 0.005

**Fig. S3.**
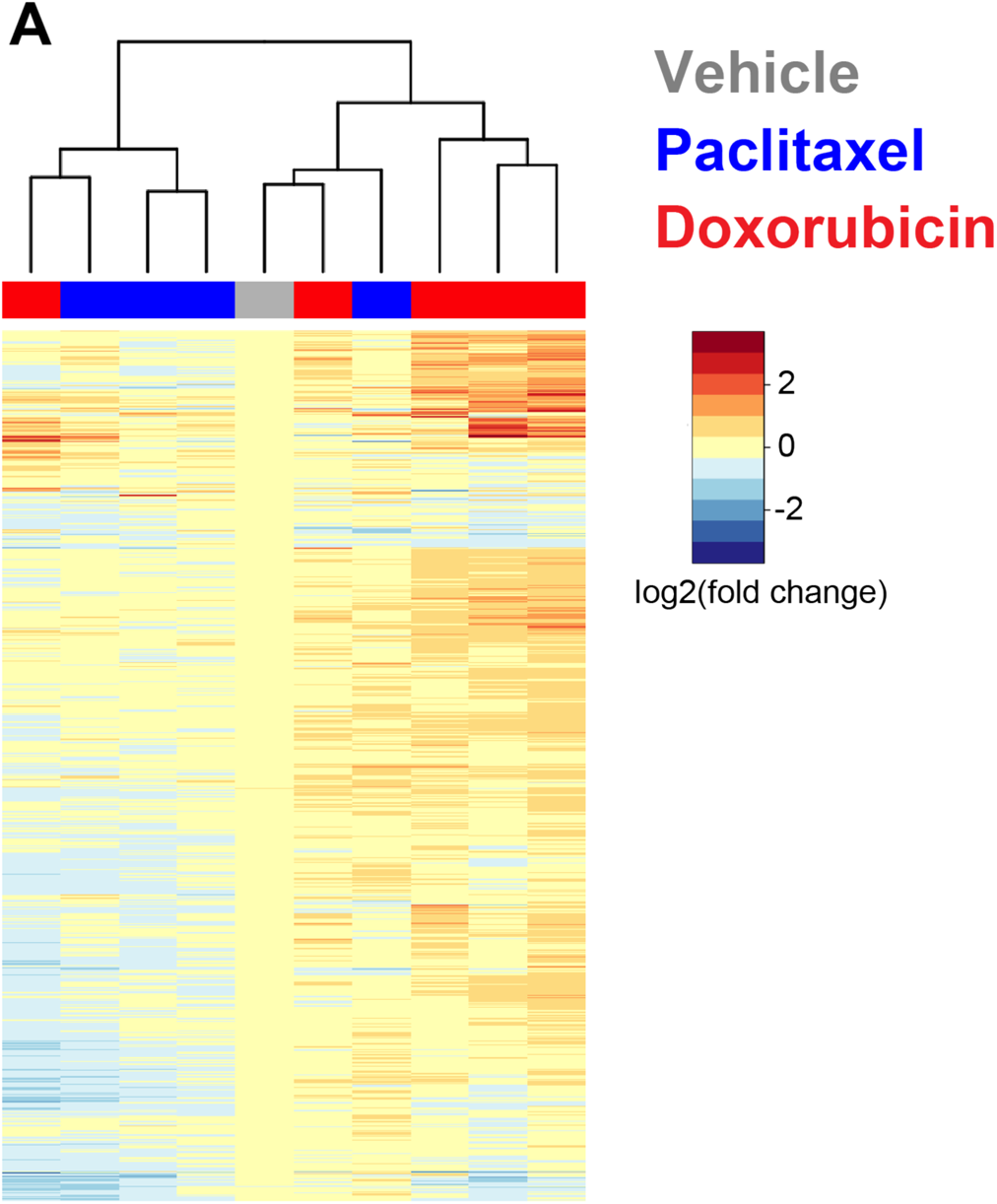
Paclitaxel and doxorubicin treatment induce largely distinct changes in tumor ECM composition. A) Heatmap and hierarchical clustering displaying results of TMT proteomics study comparing composition of dECM scaffolds from paclitaxel-, doxorubicin-, or vehicle-treated tumors.

**Fig. S4.**
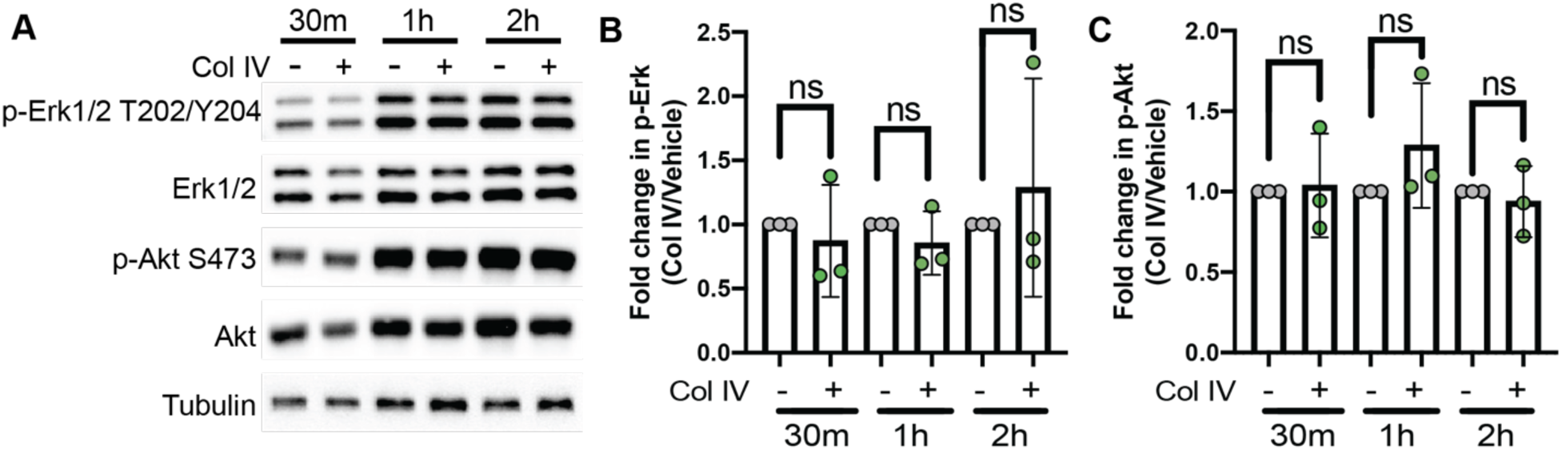
MAPK and PI3K pathways are not activated by collagen IV in PyMT-GFP TNBC cells. A) Representative western blot images of whole cell PyMT-GFP lysates immunoblotted for the indicated proteins or phosphoproteins. Cells were plated on collagen IV-coated (Col IV, 20μg/mL) or control dishes for the indicated times before lysis. B) Quantification of phospho-Erk1/2 signal as a measure of MAPK pathway activation relative to total Erk1/2 protein and normalized to a loading control. C) Quantification of phospho-Akt signal as a measure of PI3K pathway activation relative to total Akt protein and normalized to a loading control. Data show mean ± SD (n = at least 3 independent experiments). Significance was determined by unpaired *t* test.

**Table S1. Results of TMT proteomics study investigating composition of vehicle-, paclitaxel-, and doxorubicin-treated dECM.** Tumor dECM scaffolds were analyzed by LC-MS/MS and differences were quantified as log2 (Fold change) compared to control scaffolds. Data was annotated using publicly available tools (matrisome.org) to identify changes in ECM composition.

**Movie S1. PyMT-GFP cells seeded onto vehicle-, paclitaxel-, or doxorubicin-treated dECM scaffolds.** Representative videos of PyMT-GFP reseeding experiments using vehicle-(A), paclitaxel-(B), or doxorubicin-treated (C) dECM scaffolds (scale bar, 100μm).

**Movie S2. PyMT-GFP cells seeded on vehicle-treated dECM scaffolds are insensitive to Src, FAK, and integrin β1 inhibition.** Representative videos of PyMT-GFP reseeding experiments on vehicle-treated dECM with DMSO (A), dasatinib (B, 10nM), defactinib (C, 1μM), or an integrin β1 blocking antibody (D, 100ng/mL). Scale bar, 100μm.

**Movie S3. Inhibition of Src or FAK, but not integrin β1, suppresses PyMT-GFP invasion on paclitaxel-treated dECM.** Representative videos of PyMT-GFP reseeding experiments on paclitaxel-treated dECM with DMSO (A), dasatinib (B, 10nM), defactinib (C, 1μM), or an integrin β1 blocking antibody (D, 100ng/mL). Scale bar, 100μm.

**Movie S4. 231-GFP cells seeded on vehicle-treated dECM scaffolds are insensitive to Src or FAK inhibition.** Representative videos of 231-GFP reseeding experiments on vehicle-treated dECM with DMSO (A), dasatinib (B, 10nM), or defactinib (C, 1μM). Scale bar, 100μm.

**Movie S5. Inhibition of Src or FAK suppresses 231-GFP invasion on paclitaxel-treated dECM.** Representative videos of 231-GFP reseeding experiments on paclitaxel-treated dECM with DMSO (A), dasatinib (B, 10nM), or defactinib (C, 1μM). Scale bar, 100μm.

